# Early Indirect Neurogenesis transitions to late Direct Neurogenesis in mouse cerebral cortex development

**DOI:** 10.1101/2025.05.22.655488

**Authors:** Adrián Cárdenas, Irem Çelik, Alexandre Espinós, Carmen Streicher, Lara López-González, Lucia del-Valle-Anton, Virginia Fernández, Salma Amin, Enrico Negri, Eduardo Fernández Ortuño, Simon Hippenmeyer, Víctor Borrell

**Affiliations:** Instituto de Neurociencias, Consejo Superior de Investigaciones Científicas & Universidad Miguel Hernández, Sant Joan d’Alacant 03550, Spain; Institute of Science and Technology Austria (ISTA), Am Campus 1, 3400 Klosterneuburg, Austria; Current address: Department of Biological Sciences, Columbia University, New York, NY 10027, USA; Human Technopole, Milan, Italy

## Abstract

The cerebral cortex must contain the appropriate numbers of neurons in each layer to acquire its proper functional organization. Accordingly, neurogenesis requires precise regulation along development. Cortical neurons are made either directly by Radial Glia Cells (RGCs) that self- consume, or indirectly from RGCs via Intermediate Progenitor Cells (IPCs) and largely preserving the RGC pool. According to the standing model of cortical development, Direct Neurogenesis predominates at early stages of development, and progressively shifts to Indirect Neurogenesis, which predominates at late stages. However, neurogenesis at early stages should be compatible with RGC amplification, and neurogenesis at late stages needs to involve RGC consumption, which seems in conflict with the standing model. Here we studied the modes of neurogenesis along cortical development using multiple approaches, including birthdating, live imaging and MADM clone labeling. Contrary to the established dogma, our data show that Indirect Neurogenesis clearly predominates at early developmental stages, gradually shifting to Direct Neurogenesis at late stages. These findings challenge the current model of cortical neurogenesis, and prompt a re-evaluation of previous and ongoing work about the genetic and molecular mechanisms regulating this process.

## Main text

The regulation of neurogenesis during development is a critical process because it determines the types of neurons, and the numbers of each type, that will constitute the mature brain. In the developing cerebral cortex, excitatory neurons derive from the primary type of neural stem cell, apical Radial Glia Cells (aRGCs), which form the Ventricular Zone (VZ) and divide apically at the ventricular surface (Malatesta *et al*, 2000; Noctor *et al*, 2001). During neurogenesis aRGCs may produce other aRGCs, neurons or IPCs, a secondary type of neural stem cells that form the Subventricular Zone (SVZ) and divide at positions basal to the VZ (Noctor *et al*, 2004; Miyata *et al*, 2004). Neurogenesis occurring directly from aRGCs is called Direct Neurogenesis (DN), while the generation of neurons via IPCs is referred to as Indirect Neurogenesis (IDN). Importantly, because IPCs derive from aRGCs, they amplify the neuronal output of the latter, hence IDN is a more productive mode of neurogenesis than DN.

The standing model of cerebral cortex development is that DN predominates at early stages, when the less abundant neurons destined to lower layers are born. Thereafter, the numbers of IPCs gradually increase and so DN gradually shifts to IDN, which predominates at late stages when the more abundant neurons destined to upper layers are produced (Paridaen & Huttner, 2014; Jabaudon, 2017; Lui *et al*, 2011). The notion of such developmental dynamics has built over decades based on multiple observations: first, the abundance of cells expressing the transcription factor Tbr2, marker of IPCs, progressively increases during development (Noctor *et al*, 2008). Second, upper layer neurons are generated at late stages from IPCs within the SVZ (Tarabykin *et al*, 2001; Kriegstein *et al*, 2006). Moreover, IPCs are largely absent in the reptile cortex, which lacks the neuron-abundant upper layers (Cheung *et al*, 2007, 2010), whereas primates typically display an expanded SVZ that produces massive numbers of IPCs and upper layer neurons (Dehay & Kennedy, 2007; Thor, 2024).

Neuron birthdating results seem to also support the temporal sequence of early DN and late IDN in mouse cortex (Vitali *et al*, 2018). However, accumulating challenge this developmental transition from DN to IDN: basal progenitors produce neurons since the onset of neurogenesis (Haubensak *et al*, 2004). Neurons of all cortical layers, both lower and upper, derive from Tbr2^+^ cells (Kowalczyk *et al*, 2009; Vasistha *et al*, 2015; Huilgol *et al*, 2025). In the mouse rostral cortex IDN predominates at early stages (Cárdenas *et al*, 2018). Accurately defining the exact temporal sequence of neurogenic events in the developing cerebral cortex is fundamental to understand its cellular and molecular regulation, and the interpretation of past and future studies.

To specifically study the modes of neurogenesis along cortical development, we first analyzed the abundance of Tbr2^+^ cells in the mouse cerebral cortex during the neurogenic period, between embryonic days (E) E12.5 and E16.5. We observed a progressive increase in Tbr2^+^ cells along development (Fig. 1a) as previously reported (Noctor *et al*, 2008). This feature was paralleled by the inside-out temporal gradient of neurogenesis in the cerebral cortex (Extended Data Fig. 1a,b) (Rakic, 1974), supporting the notion of a developmental increase in IDN. Next, we analyzed the occurrence of mitoses in apical (aRGCs) and basal positions (IPCs), identified by immunostains against Phosphorylated histone H3 (PH3). At all neurogenic stages we observed both apical and basal mitoses (Fig. 1b), again in agreement with previous reports (Noctor *et al*, 2008; Kowalczyk *et al*, 2009). However, the density of basal mitoses did not increase between E12.5 and E16.5 but remained similar, in contrast to the increase in Tbr2^+^ cells (Fig. 1c). On the other hand, the density of apical mitoses decreased to half during this period, so *de facto* the relative abundance of basal mitoses increased nearly 2-fold (Fig. 1c), potentially compatible with a developmental increase in IDN.

**Fig. 1.**
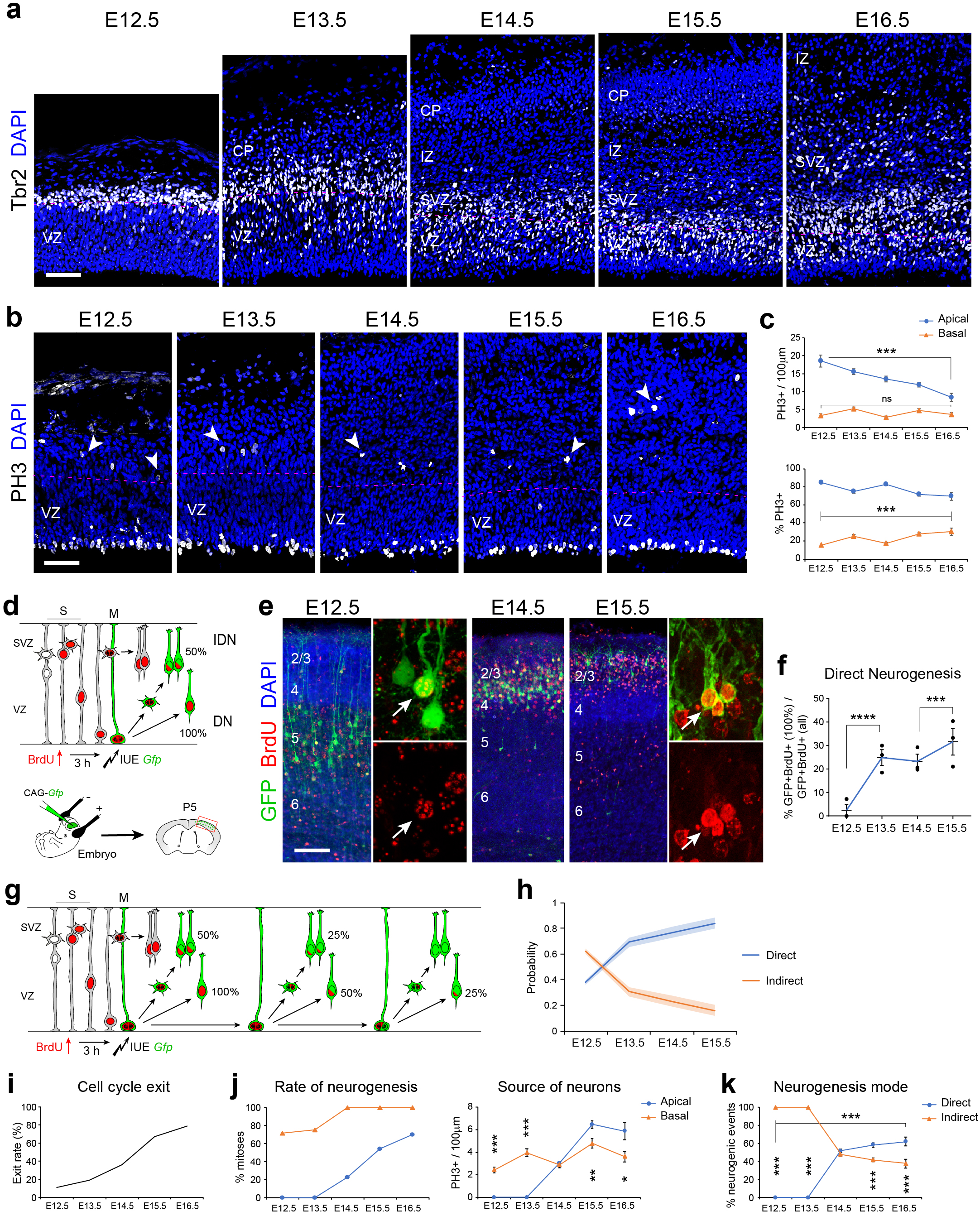
Mitotic activity and neuron birthdating show a developmental increase of Direct Neurogenesis. **a**,**b**, Coronal sections of mouse cerebral cortex at the indicated embryonic stages (E) stained for Tbr2 (a) or PH3 (b). Dotted lines indicate basal border of the ventricular zone (VZ). Arrowheads indicate mitoses at basal positions. CP, cortical plate; IZ, intermediate zone; SVZ, subventricular zone. **c**, Quantification of density and relative abundance of apical and basal PH3+ cells as indicated (*n* = 9-17 embryos per age). **d**, Experimental design to identify neurons born by DN or IDN based on GFP labeling and BrdU retention. **e,f**, Patterns of GFP and BrdU labeling at P5 after electroporation and injection at the indicated ages (e), and quantifications (f) (*n* = 3 embryos per age; χ^2^-test). Arrows indicate positive pyramidal neurons; cortical layers are indicated by numbers. **g,h**, Schema of BrdU dilution in subsequent cell divisions leading to underestimation of DN at first division (g), and corrected probability of DN and IDN along corticogenesis (h); values are max-to-min range (light color) and mean (dark line). **i**, Cell cycle exit rate in mouse neocortex, from (Takahashi *et al*, 1996). **j**, Rate of neurogenesis from apical and basal mitoses, and density of neuron-producing mitoses, calculated for each developmental stage based on data in (c,i) (*n* = 9-17 embryos per age; t-test). **k**, Proportion of DN and IDN based on data in (j) (χ^2^-test). DN increased significantly from E12.5 to E15.5. Values are mean ± SEM; \**p* < 0.05; \*\**p* < 0.01; \*\*\**p* < 0.001; \*\*\*\**p* < 0.0001. Scale bars: 50µm (a,b), 200µm (e).

To directly determine the modes of neurogenesis occurring along cortical development, we next labeled neurons born via DN by combining single-pulse BrdU incorporation with *in utero* electroporation of *Gfp*-encoding plasmids to label apical progenitors (essentially aRGCs; Fig. 1d) (Cárdenas *et al*, 2018). Apical progenitors were labeled at different embryonic stages and all animals were analyzed at P5. GFP-labeled pyramidal neurons containing 100% BrdU were considered born by DN at the stage of BrdU injection, with lower doses of BrdU indicating additional cell divisions before neurogenesis, such as IDN (Fig. 1d; Extended Data Fig. 1c). We observed a nearly absent occurrence of DN at E12.5, which gradually increased along development to peak at E15.5 (Fig. 1e,f). An important limitation of these experiments was that BrdU labeling is carried over in self-renewed progenitors, so that we identified as IDN neurons born from IPCs and those born from aRGCs after several rounds of self-renewal. Accordingly, DN was underestimated, especially at early stages with many cell cycles still to occur (Fig. 1g). Taking into account this limitation, we used our observations on low BrdU labeling across ages to recalculate the actual frequency of DN at each age (see Methods). Our results confirmed the transition from early IDN to late DN (Fig. 1h). These results strongly supported that IDN predominates at early stages, and DN increases gradually at later stages.

In light of the above findings, we re-examined our results from PH3 stains. Pioneer studies showed that the rate of cell cycle exit along mouse corticogenesis escalates from E12.5 to E16.5 (Fig. 1i) (Takahashi *et al*, 1996). This means that at early stages very few of all mitoses are neurogenic, but mostly produce cycling progenitor cells, while at late stages the vast majority of mitoses are neurogenic. Considering that IPCs (basal mitoses) in mouse are mostly neurogenic rather than self-renewing (Noctor *et al*, 2004), the observed abundance of basal mitoses at early stages would suffice to account for the very low levels of neurogenesis, with null contribution from apical mitoses (Fig. 1j). In contrast, at late stages the moderate abundance of basal mitoses is insufficient to account for the high rate of neurogenesis (cell cycle exit), thus necessarily involving apical mitoses – up to 70% by E16.5 (Fig. 1j). Accordingly, apical progenitors produced very low numbers of neurons at early stages, but this increased during development to, eventually, nearly double the rate of neurogenesis of basal progenitors by E16.5 (Fig. 1k). Importantly, this dynamics did not change even if considering that IPCs may self-renew (Extended Data Fig. 1d). These results strongly supported that IDN is the main mode of cortical neurogenesis at early stages, and DN gradually gains relevance to eventually predominate at late stages (Fig. 1k).

To directly visualize the occurrence of DN and IDN, we monitored the fate of cortical progenitor cells by videomicroscopy. Individual aRGCs were labeled with GFP by *in utero* electroporation, and one day later the lineages of labeled progenitor cells were monitored by time-lapse imaging in brain slices, followed by cell type-specific marker analysis: Pax6 for aRGCs, Tbr2 for IPCs, Neurod1 for neurons (Fig. 2a) (Cárdenas *et al*, 2018). At E12.5, aRGCs produced essentially aRGCs (79%) or IPCs (18%), while neuron production from aRGCs (DN) was minimal (3%; Fig. 2b,c,f,g; Extended Data Fig. 2; Extended Data Movies 1-5). At this early age, most neurons were born from IPCs (89%; Fig. 2f,g), altogether indicating the predominance of IDN. In contrast, at E15.5 aRGCs produced much fewer aRGCs (50%) but more IPCs (38%) and, especially, neurons (12%; Fig. 2d-g; Extended Data Fig. 3,4; Extended Data Movies 6-11). Indeed, the rate of DN increased 4-fold between E12.5 and E15.5 (Fig. 2g), again revealing a developmental increase in DN.

**Fig. 2.**
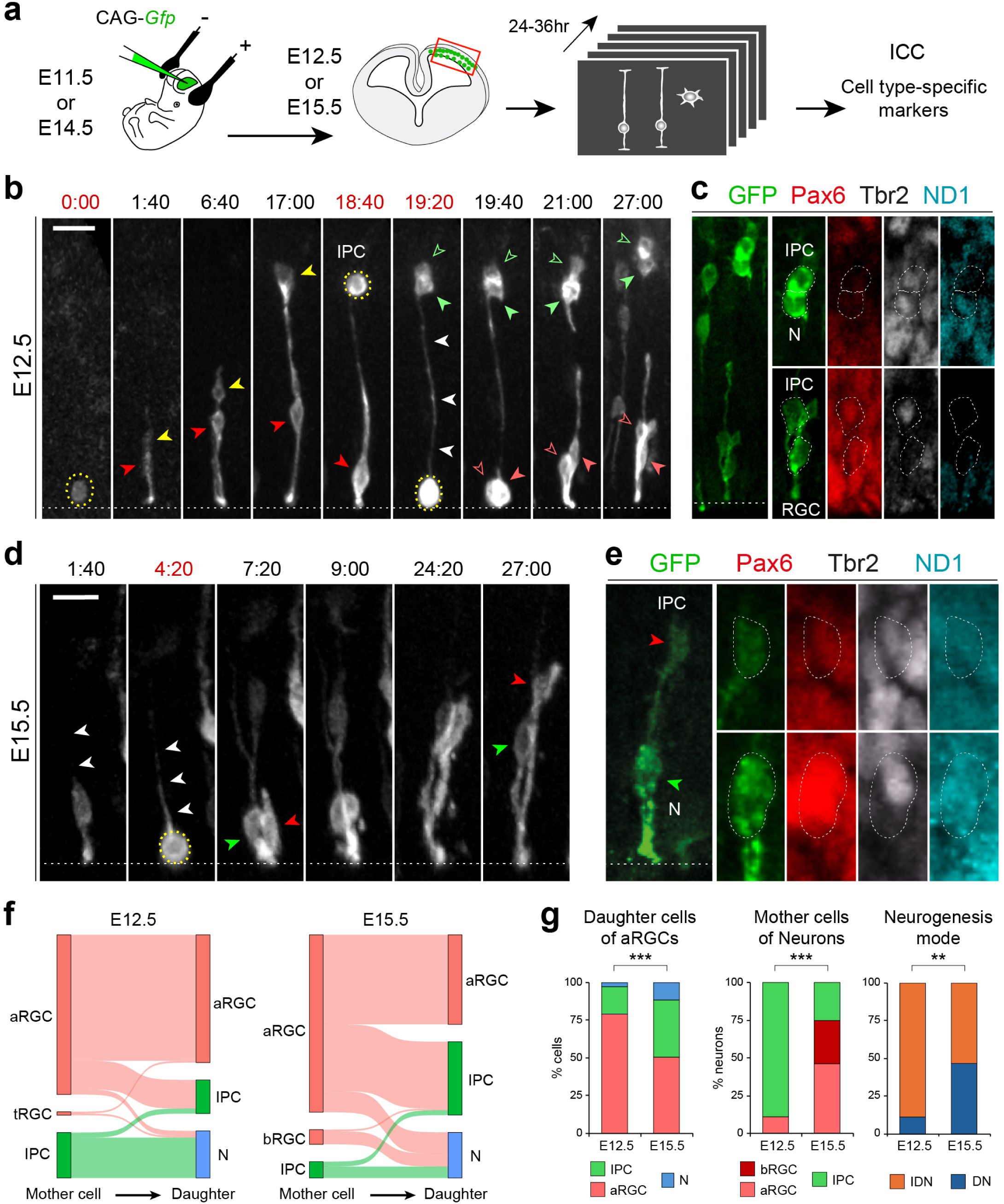
Live imaging reveals a developmental shift from IDN to DN. **a**, Experimental design of time-lapse imaging: embryos are electroporated *in utero* with CAG-*Gfp*-encoding plasmids, one day later living brain slices are obtained and labeled progenitor cells are imaged for 24-36hrs. After imaging, tissue slices are fixed and stained for cell type-specific markers. **b-e**, Time-lapse imaging frames from embryos at the indicated ages (b,d), and post-hoc marker staining (c,e), showing examples of RGCs undergoing IDN (E12.5) and DN (E15.5). Stains for GFP, Pax6, Tbr2 and NeuroD1 (ND1) after each movie are shown, with the identity of daughter cells indicated. White arrowheads indicate basal process of aRGCs; yellow dashed circles indicate mitotic cells; red, yellow and green arrowheads indicate daughter RGCs, IPCs and neurons (N), respectively; white dotted lines indicate apical VZ surface. Time stamps are in hr:min; red indicates time at mitosis. Scale bars: 15µm. **f,** Sankey diagrams of cortical clones identified by live imaging, with confirmed daughter cell identities. aRGC, apical Radial Glia Cell; bRGC, basal Radial Glia Cell; tRGC, truncated Radial Glia Cell; IPC, Intermediate Progenitor Cell; N, neuron (*n* = 76 clones E12.5; 86 clones E15.5). **g**, Quantifications of live-imaging clone analyses indicating daughter cells produced by aRGCs (*n* = 95 cells E12.5; 109 cells E15.5), neurons born from each progenitor type and neurogenesis mode (*n* = 27 neurons E12.5; 28 neurons E15.5). \*\**p*<0.01; \*\*\**p*<0.001, χ^2^-test. Scale bars: 15µm.

A caveat of these experiments was that, whereas the cells originally labeled by *in utero* electroporation were apical progenitors, some of these had divided already before the start of imaging, generating aRGCs, IPCs and neurons (Extended Data Figs. 2-4). Thus, the frequency in which we imaged neurons being born from aRGCs or IPCs was biased by the relative abundance of aRGCs and IPCs labeled at the start of imaging, and monitored thereafter. At E12.5 we observed few aRGCs producing IPCs (18% daughter cells; Fig. 2g). Despite this, most neurons were born from IPCs at this age (89%; Fig. 2f,g). Conversely, at E15.5 aRGCs produced 2-fold more IPCs than at E12.5 (38% of daughter cells), and yet the frequency of neurons being born from IPCs diminished to 25% of total neurons - perhaps owing to increased cell cycle length in IPCs - while aRGCs produced 46% of neurons (Fig. 2g). Altogether, this strongly supported a predominance of IDN at E12.5 and a dramatic increase of DN by E15.5.

We next aimed to corroborate our above findings concerning the modes of cortical neurogenesis *in vivo* with a genetic strategy, using the Mosaic Analysis with Double Markers (MADM) technology (Zong *et al*, 2005; Gao *et al*, 2014; Contreras *et al*, 2021). We used a tamoxifen (TM)-inducible *Emx1*-CreER recombinase driver line to induce MADM clones from cortical aRGCs starting at specific stages of embryonic development, and animals were analyzed at maturity (P28) (Gao *et al*, 2014; Beattie *et al*, 2020). We focused only on two color MADM clones, containing green and red cells, and analyzed the composition of their individual sub-clones (green or red; referred to as clones hereafter). We reasoned that clones containing only one neuron in one color reflected DN, and those with more than one neuron reflected non-terminal neurogenesis at the time of labeling (Fig. 3a). Overall, MADM clones induced at early stages contained neurons in all layers, and those induced at gradually later stages became progressively enriched in more superficial layers, as expected given the inside-out gradient of neurogenesis (Fig. 3b-d). At the level of individual clones, a majority of clones induced at each age contained neurons located in the layers corresponding to that birthdate (i.e. most E12.5-induced clones contained neurons in layers 6 and 5; most E14.5 clones contained neurons in layers 5 and 4), plus neurons in more superficial layers due to the persistent labeling of the mother cell (Fig. 3e). To focus on neurogenic events taking place specifically at the embryonic stage of clone induction, we analyzed only clones that contained neurons in the cortical layers corresponding to that birthdate (i.e. clones induced at E12.5 containing neurons in layers 6 or 5). Clones without neurons in those layers likely resulted from an initial self-amplification of aRGCs or IPCs, thereby producing neurons only at later stages and thus destined only to more superficial layers. Our analysis revealed that the frequency of DN clones was only 18% at E12.5, gradually increasing to 56% at E15.5 (Fig. 3f). Two-color MADM clones containing a single neuron of each color (DN in both clones) also increased significantly from E12.5 to E15.5 (Fig. 3g). Altogether, this confirmed the gradual transition from IDN to DN along mouse cerebral cortex development.

**Fig. 3.**
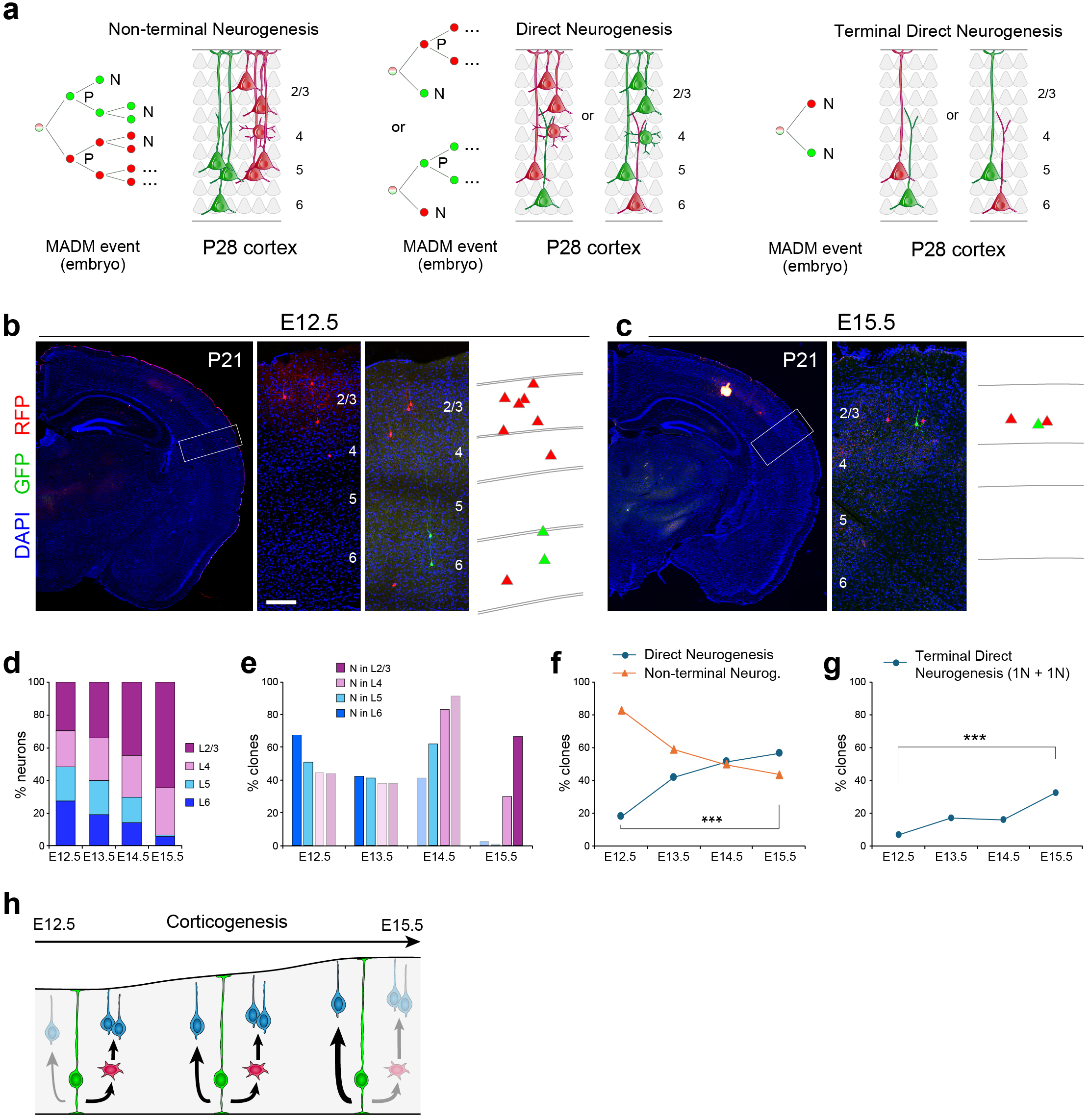
MADM clonal analysis corroborates a developmental shift from IDN to DN. **a**, Schemes of MADM clone architectures: left, Non-terminal Neurogenesis; middle, Direct Neurogenesis; right, Terminal Direct Neurogenesis. N, neuron; P, progenitor. **b,c**, Representative images of brain sections from *MADM-11^GT/TG^;Emx1-CreERT2^+/-^*mice at P28, indicating MADM clones induced at E12.5 (b) and E15.5 (c), respectively; and reconstruction of green- and red-labeled clones. Boxes indicate areas shown at higher magnification. **d,e**, Frequency of laminar distribution of MADM- labeled neurons from all clones, induced at respective embryonic ages (E12.5, E13.5, E14.5 and E15.5) (d), and proportion of clones induced at each age that contain neurons in each layer (e) (*n* = 66-140 clones, 7-22 brains per induction time-point). Most clones have neurons in two layers (dark colors) corresponding to the location of those born circa the stage of MADM induction. This subset of clones reveals the type of neurogenesis at each stage. **f,g**, Proportion of clones indicating Direct Neurogenesis and Non-terminal neurogenesis along development (f), and proportion of two color MADM clones indicating terminal Direct neurogenesis (g), as indicated (*n* = 60-123 clones per induction time-point; \*\*\**p*<0.001, χ^2^-test). **h**, Summary schema: IDN predominates at early stages, but remains similar along development while DN increases progressively, to predominate at late stages. Scale bars: 100µm.

Our results demonstrate that IDN is the predominant mode of neurogenesis at the onset of cortical development. IDN frequency decreases progressively during the neurogenic period concomitant with an increase in DN, which becomes predominant at the last stages of cortical neurogenesis (Fig. 3h). These results contrast with the widely accepted model of cortical neurogenesis, according to which DN predominates at early stages and IDN gradually becomes predominant as development progresses (Paridaen & Huttner, 2014; Jabaudon, 2017; Lui *et al*, 2011). Whereas the coexistence of DN and IDN all along development is widely recognized, the established model stands largely based on two key observations: the developmental increase in Tbr2^+^ IPCs and in the thickness of SVZ where they reside. This model is also coherent with the increased numbers of neurons generated at late developmental stages and destined to form upper layers. However, progenitor cells must divide to produce neurons, and the duration of the cell cycle increases very significantly as development proceeds, such that larger populations of IPCs dividing more slowly may result in similar amounts of neurons produced during a given developmental time. The similar abundance of basal mitoses observed across development supports this notion. The standing model seems to also neglect two other key parameters when considering DN vs IDN: cell cycle exit rate, and progenitor cell consumption. At early developmental stages, progenitor cells must undergo self-amplifying divisions, such that the rate of cell cycle exit is very low. Under these conditions, even a small pool of IPCs, which are largely self-consumed when producing neurons, is sufficient to account for all the neurogenic output without the involvement of RGCs (DN). IDN predominates at early stages even under the assumption that 50% of IPCs self-renew (Extended Data Fig. 1d), which in mouse cortex is rare (Noctor *et al*, 2008, 2004). In contrast, at late stages of neurogenesis the observed abundance of basal (IPC) mitoses alone is insufficient to account for the very high rates of cell cycle exit (Takahashi *et al*, 1996), requiring the involvement of a large number of apical (RGC) mitoses and hence a shift toward DN predominance. Importantly, during the last period of cortical neurogenesis before RGCs produce or become astrocytes, VZ thickness and RGC abundance decrease dramatically, indicating high RGC self-consumption and consistent with high rates of DN.

Recent lineage-tracing analyses based on TM-inducible *Tbr2*-CreER based fate mapping reported predominant DN in the cerebral cortex at early stages producing deep layer neurons, and a developmental increase in IDN significantly contributing to upper layer production (Huilgol *et al*, 2023). While these inducible transgenic lines may be of great value to study cell fate, their validation revealed only partial labeling of RGCs and IPCs. Potentially due to an incomplete effect of tamoxifen, or other unexplored options, the partial labeling warns about the fundamental limitations of this approach in the quantitative assessment of DN vs IDN. On the other hand, an independent study using a similar tamoxifen-inducible strategy, but under the control of a different reporter line, recently showed high levels of IDN at early stages of cortical development (Hatanaka *et al*, 2024), in agreement with the findings in this present study.

Our findings in mouse prompt the question of phylogenic conservation in the developmental shift from IDN at early stages to DN at late stages. Previous studies show that in ferret, with an SVZ expanded and split into inner and outer SVZ (ISVZ, OSVZ), progenitor cells in the OSVZ follow a lineage independent from VZ and ISVZ, with massive self-amplification at mid-stages of neurogenesis (Martínez-Martínez *et al*, 2016), suggesting a completely different scenario. Similarly, cortical neurogenesis in macaque monkeys slows down at mid stages, when progenitors self-amplify before undertaking the production of massive amounts of upper-layer neurons in the latest period (Betizeau *et al*, 2013). As for neurogenesis from VZ-ISVZ, recent analyses show evidence of DN and IDN in ferret (Del-Valle-Anton *et al*, 2024; Singh *et al*, 2024). These also reveal multiple neurogenic lineages occurring simultaneously and in parallel in ferret and human cortex, but not mouse (Del-Valle-Anton *et al*, 2024), strongly suggesting a much more complex scenario in species with a larger and more complex cortex. The dynamics of DN and IDN in large-brained species, including human, remains a fascinating open question for future research.

## Author contributions

V.B. conceived the project and designed the experiments; A.C. performed and analyzed videomicroscopy experiments; C.S. performed MADM experiments, analyzed by I.C. with the guidance and supervision of S.H. and V.B.; A.C., E.F., A.E., P.A. and V.B. performed and analyzed mitotic labeling and birthdating experiments; E.N. modeled birthdating and neurogenesis dynamics; L.L.-G., V.F., L.dV.A. and S.A. performed scRNA-seq experiments, analyzed by L.L.-G. and L.dV.A.; S.H. and V.B. provided reagents and resources. V.B. supervised experiments, provided funding, prepared figures and wrote the manuscript with input from all other authors.

## Acknowledgements

We thank A. Iñigo for assistance with imaging, and members of the Borrell and Herrera labs for insightful discussions and critical reading of the manuscript. Funding to our lab members was provided by the Spanish Research Agency (AEI): FPI contract (BES-2016-077737) to L.dV.A., FPI SO contract (SEV-2017-0723-18-1) to A.E., JdC-Incorporación contract (IJC2020-044653-I) to V.F., and JAE-Intro fellowship (JAEICU23EX_0071) to I.C., as well as by La Caixa Foundation: La Caixa-Severo Ochoa fellowship (E-03-2016-0557140) to S.A., INPhINIT-Retaining fellowship (LCF/BQ/DR21/11880012) to E.F.O., INPhINIT-Incoming fellowship (LCF/BQ/DI22/11940006) to E.N., and Junior Leader-Retaining grant to A.C. (LCF/BQ/PR23/11980051). Work was supported by grants from FWF (SFB F78) to S.H.; AEI (PID2021-125618NB-I00) and European Research Council (101118729) to V.B., who also acknowledges financial support from AEI through the “Severo Ochoa” Programme for Centers of Excellence in R&D (CEX2021-001165-S).

## Extended Data Figure legends

**Extended Data** Fig. 1**. Layering of cortical neurons during development follows an inside-out pattern. a**, Coronal sections of mouse neocortex at P5 showing the patterns of BrdU label (white) after a single dose injected at the indicated embryonic stages. **b**, Binned quantification of BrdU density across the cortical thickness (a.u., normalized arbitrary units; *n* = 3 pups per age; mean values are shown). **c**, Representative examples of cortical pyramidal neurons at P21 expressing GFP after electroporation at embryonic stages, and showing the indicated levels of BrdU retention. Insets detail the amount and pattern of nuclear BrdU staining. **d**, Rate of neurogenesis from Apical and Basal mitoses, density of neuron-producing mitoses along development, and proportion of DN and IDN, depending on the cell cycle exit rate of IPCs (color code). Results at each developmental stage are based on data in main Fig 1c,i. In all conditions, IDN decreased and DN increased between E12.5 and E15.5. Scale bars = 100mm (a), 15mm (c).

**Extended Data** Fig. 2**. Examples of time-lapse movies at E12.5.** Individual frames from time-lapse imaging (time stamps are indicated in hr:min of imaging), and stains for GFP and cell type-specific markers (Pax6, Tbr2, NeuroD1 - ND1) of imaged tissue after the last frame. The identity of daughter cells is indicated. **a,b**, Examples of aRGC self-amplification. **c**, Example of Direct Neurogenesis. **d**, Example of Indirect Neurogenesis; the first mitosis generates one RGC (red arrowhead) that goes out of focus after 18:00hr. White arrowheads indicate initial aRGCs; yellow dashed circles indicate mitotic cells; red, yellow and green arrowheads indicate daughter RGCs, IPCs and neurons (N), respectively; white dotted lines indicate apical VZ surface. Time stamps in red indicate time at mitosis. Scale bars = 30mm.

**Extended Data** Fig. 3**. Examples of time-lapse movies at E15.5.** Individual frames from time-lapse imaging (time stamps are indicated in hr:min of imaging), and stains for GFP and cell type-specific markers (Pax6, Tbr2, NeuroD1 - ND1) of imaged tissue after the last frame. The identity of daughter cells is indicated. **a,b**, Examples of aRGC self-amplification. **c**, Example of generation of IPC. **d**, Example of Direct Neurogenesis. **e**, Example of Direct Neurogenesis plus generation of IPC. White arrowheads indicate initial aRGCs; yellow dashed circles indicate mitotic cells; red, yellow and green arrowheads indicate daughter RGCs, IPCs and neurons (N), respectively; white dotted lines indicate apical VZ surface. Time stamps in red indicate time at mitosis. Scale bars = 30mm.

**Extended Data** Fig. 4**. Sankey diagrams of E12.5 and E15.5 cortical lineages analyzed by time-lapse imaging. a**, Analysis of lineages based on cell morphology (*n* = 260 lineages at E12.5; 159 at E15.5). **b**, Analysis of lineages based on marker expression (*n* = 54 lineages at E12.5; 65 at E15.5). aRGC, apical Radial Glia Cell; bRGC, basal Radial Glia Cell; tRGC, truncated Radial Glia Cell; BP, bipolar cell; saP, subapical Progenitor; IPC, Intermediate Progenitor Cell; abIPC, apico-basal process IPC; MP, multipolar cell; N, neuron; ND1, Neurod1.

**Extended Data Movie 1. Indirect Neurogenesis at E12.5.** Videomicroscopy of the lineage of a single aRGC in an organotypic slice culture from the mouse neocortex at E12.5, as shown in Fig. 2b. Apical surface is down. The slice was prepared 24hr after *in utero* electroporation, and imaging started 4hrs after slice preparation. Total imaging time is 27hrs. Colored arrowheads follow the same RGC and its progeny through the movie. The initial RGC divides at the apical surface at 0:40hr and produces 1 IPC (yellow arrowhead) plus 1 RGC (red arrowhead); the RGC undergoes INM again to divide at the apical surface and produce 2 more RGCs (open red arrowheads), whereas the IPC moves basally to undergo a terminal division without INM, producing 2 neurons (green arrowheads).

**Extended Data Movie 2. RGC amplification at E12.5.** Videomicroscopy of the lineage of a single aRGC in an organotypic slice culture from the mouse neocortex at E12.5, as shown in Extended Data Fig. 2a. Apical surface is down. The initial RGC (white arrowhead) undergoes interkinetic nuclear migration to divide apically and produce 2 RGCs (red arrowheads).

**Extended Data Movie 3. RGC amplification at E12.5.** Videomicroscopy of the lineage of a single aRGC at E12.5, as shown in Extended Data Fig. 2b. Apical surface is down. The initial RGC (white arrowhead) undergoes interkinetic nuclear migration to divide apically and produce 2 RGCs (red arrowheads). The initial RGC undergoes interkinetic nuclear migration (INM) to divide apically and produce 2 RGCs (red arrowheads), which undergo INM again and one of them divides at the apical surface to produce 2 more RGCs, for a total of 3 at the end of imaging time.

**Extended Data Movie 4. Direct Neurogenesis at E12.5.** Videomicroscopy of the lineage of an aRGC at E12.5, as shown in Extended Data Fig. 2c. Apical surface is down. The initial RGC (white arrowhead) divides apically to produce 1 RGC (red arrowhead) plus 1 N (green arrowhead). Then the RGC undergoes interkinetic nuclear migration to divide apically again and produce 2 RGCs.

**Extended Data Movie 5. Indirect Neurogenesis at E12.5.** Videomicroscopy of the lineage of an aRGC at E12.5, as shown in Extended Data Fig. 2d. Apical surface is down. The initial RGC (white arrowhead) divides apically to produce 1 RGC (red arrowhead) plus 1 IPC (yellow arrowhead). Both cells move to the basal side, the RGC goes out of focus, and the IPC divides basally to produce 2 neurons (green arrowheads).

**Extended Data Movie 6. Direct Neurogenesis at E15.5.** Videomicroscopy of the lineage of a single aRGC in an organotypic slice culture from mouse neocortex at E15.5, as shown in Fig. 2d. Apical surface is down. Total imaging time is 27hrs. Colored arrowheads follow the same RGC and its progeny through the movie. The initial RGC divides at the apical surface to produce 2 neurons (green arrowheads) that remain attached to the apical surface until the end of the movie. These cells were identified as neurons based on *post-hoc* marker expression analysis.

**Extended Data Movie 7. RGC amplification at E15.5.** Videomicroscopy of the lineage of a single aRGC at E15.5, as shown in Extended Data Fig. 3a. Apical surface is down. The initial RGC (white arrowhead) undergoes interkinetic nuclear migration to divide apically and produce 2 RGCs (red arrowheads).

**Extended Data Movie 8. RGC amplification at E15.5.** Videomicroscopy of the lineage of two individual aRGCs at E15.5, as shown in Extended Data Fig. 3b. Apical surface is down. The initial RGCs (white arrowheads) undergo interkinetic nuclear migration to divide apically and produce 2 RGCs each (red arrowheads).

**Extended Data Movie 9. Generation of IPC at E15.5.** Videomicroscopy of the lineage of a single aRGC in an organotypic slice culture from mouse neocortex at E15.5, as shown in Extended Data Fig. 3c. Apical surface is down. The initial RGC (white arrowhead) undergoes interkinetic nuclear migration to divide apically and produce 1 RGC (yellow arrowhead) plus 1 IPC (red arrowhead), which remains attached apically until the end of the movie and was identified based on *post-hoc* marker expression analysis.

**Extended Data Movie 10. Direct Neurogenesis at E15.5.** Videomicroscopy of the lineage of a single aRGC from mouse neocortex at E15.5, as shown in Extended Data Fig. 3d. Apical surface is down. The initial RGC (white arrowhead) divides at the apical surface to produce 1 RGC (red arrowhead) plus 1 neuron (green arrowhead). Cell identity was confirmed by *post-hoc* marker expression analysis.

**Extended Data Movie 11. Direct Neurogenesis plus IPC-genesis at E15.5.** Videomicroscopy of the lineage of a single aRGC from mouse neocortex at E15.5, as shown in Extended Data Fig. 3e. Apical surface is down. The initial RGC (white arrowhead) divides at the apical surface to produce 1 IPC (yellow arrowhead) plus 1 neuron (green arrowhead). Cell identity was confirmed by *post-hoc* marker expression analysis.

## Materials and Methods

### Animals

Wild-type mice were maintained in an ICR background and bred at the Instituto de Neurociencias de Alicante in accordance with Spanish and EU regulations. The day of vaginal plug was considered as embryonic day (E) 0.5. For MADM experiments, all animal procedures were approved by the Austrian Federal Ministry of Science and Research in accordance with the Austrian and European Union animal law (license number: BMWF-66.018/0007-II/3b/2012 and BMWFW- 66.018/0006-WF/V/3b/2017). Experimental mice were bred and maintained according to regulations approved by institutional animal care and use committee, institutional ethics committee and the guidelines of the preclinical facility (PCF) at ISTA. Transgenic mouse lines with MADM reporter cassettes inserted in Chr.11 (Hippenmeyer *et al*, 2010) and *Emx1*-CreER driver (Kessaris *et al*, 2006) have been previously described. For MADM experiments, mouse lines were kept in mixed C57/Bl6, FVB and CD1 genetic background. In all experiments, both male and female littermates of the desired genotypes were used randomly. Mice were used at an age range from 2-8 months for breeding and from E12.5 to P28 for experiments. All efforts were made to minimize the number of animals used following the 3R principles.

### BrdU labeling

Bromodeoxyuridine (BrdU, SIGMA) was diluted at 10mg/ml in 0.9% NaCl and always administered at 50mg/kg body weight. For birthdating analysis, a single dose of BrdU was intraperitoneally injected in timed-pregnant mice at E12.5, E13.5, E14.5, E15.5 or E16.5, and the brains of labeled animals were analyzed at postnatal day (P) P5. For experiments combining BrdU dilution with *in utero* electroporation, pregnant mice were injected at E12.5, E13.5, E14.5 or E15.5 with a single pulse of BrdU, 3hr later embryos were electroporated in the NCx with CAG-GFP plasmids, and analyzed at P21. Animals were perfused with 4% paraformaldehyde (PFA) in 0.1M phosphate buffer (PB) pH7.3, and their brains were postfixed for 1hr in ice-cold 4% PFA before processing for IHC.

### *In utero* electroporation

Mouse embryos were electroporated *in utero* in the NCx as described (Cárdenas *et al*, 2018). Briefly, pregnant females were deeply anesthetized with isoflurane and the uterine horns exposed; DNA solution (1-2µl) was injected into the lateral ventricle using pulled glass micropipettes, and square electric pulses (30-45V, 50ms on – 950ms off, 5 pulses) were applied with an electric stimulator (Cuy21EDIT Bex C., LTD) using round electrodes (CUY650P5, Nepa Gene). For birthdating experiments, pCAG-EGFP-encoding plasmids were injected at 1µg/µl.

### Immunohistochemistry

Pregnant mice were euthanized by cervical dislocation, and postnatal animals were deeply anesthetized with a mixture of ketamine (65mg/kg), xylazine (13mg/kg) and acepromazine (2mg/kg) by IP injection. Brains were fixed with 4% paraformaldehyde (PFA) in phosphate buffer (PB) pH7.3 at 4°C, either by transcardiac perfusion (postnatal animals) or by immersion following decapitation (embryos). Embryonic brains were cryoprotected with 30% sucrose, embedded in Cryo-medium Neg-50 (Thermo Scientific), frozen and cryosectioned. Postnatal brains were embedded in agarose and vibratome-sectioned. Brain sections were permeabilized in PBS containing 0.25% Triton X-100 and blocked in 10% of Horse Serum and 2% Bovine Serum Albumin (BSA) during 2 hours; then, sections were incubated with primary antibodies overnight, followed by appropriate fluorophore-conjugated secondary antibodies and counterstained with DAPI. For BrdU detection, sections were fist subject to antigen retrieval with 2N HCl at 37°C for 30 min. Primary antibodies used were: anti-BrdU (1:200, rat monoclonal, Abcam); anti-GFP (1:1000, chicken polyclonal, Aves Lab.); anti-phosphohistone H3 (1:500, mouse monoclonal, Millipore); anti-Tbr2 (1:250, rabbit polyclonal, Abcam); anti-Pax6 (1:500, rat monoclonal, Biolegend) and anti-NeuroD1 (1:250, goat polyclonal, Invitrogen). Secondary antibodies used were: biotinylated anti-Rabbit IgG and biotinylated anti-Goat IgG (Vector); Alexa488, Alexa555 and Alexa647 donkey anti-mouse and anti-rabbit IgG (Invitrogen); Cy3 donkey anti-rat, Alexa488 donkey anti-chicken IgY, DvLight405, Cy2- and Cy5-streptavidin (Jackson Immunoresearch); all diluted 1:200.

### Live imaging of progenitor cells

Mouse embryos were electroporated *in utero* at E11.5 or E14.5 with pCAG-Flox-farnesylated-EGFP (0,4µg/µl) plus pCAG-Cre (10ng/µl) plasmids. The low concentration of Cre plasmid allowed sparse labelling of individual apical progenitor cells upon recombination of the floxed stop cassette (Cárdenas *et al*, 2018). One day after electroporation, the live brains were dissected out and vibratome sliced at 275μm in ice-cold DMEM-F12 (Gibco) supplemented with glucose (2,9g/l), sodium bicarbonate (1,2g/l) and PenStrep (1ooU/ml), bubbled with carbogen (5% carbon dioxide + 95% oxygen). Slices were embedded in collagen matrix (Nitta gelatin) on a filter membrane (Millipore) and maintained in HEPES-buffered DMEM-F12 (Gibco), 5% fetal bovine serum, 5% horse serum, N2 (1:100; Invitrogen), B27 (1:50; Invitrogen) and PenStrep (100U/ml) (Cárdenas *et al*, 2018). Imaging was performed on an inverted spinning-disk microscope (Andor Dragonfly Spinning Disk Confocal microscope), 25x immersion lens and 5%CO2/37°C atmosphere. Z-stacks of frames separated 4µm were captured every 20min for 24-36hr. Immediately after recording the last time-point, slices were fixed in 4% PFA for 10min and processed for IHC. Digital images were acquired, contrast-enhanced and analyzed with Imaris software (Bitplane) and Fiji (ImageJ). Each individual mitosis was followed until the end of the movie to determine cell types derived from it, to then define the cell lineage. For co-localization analyses, images from a single confocal plane were obtained using an Olympus FV10 confocal microscope.

### MADM experiments

MADM clone induction in the telencephalon was adapted from previously described protocols (Hippenmeyer et al, 2010; Gao et al, 2014; Beattie et al, 2020). In brief, *MADM- 11^GT/TG^;Emx1-CreER^+/-^* experimental mice were generated by crossing *MADM-11^GT/GT^;Emx1- CreER^+/-^*with *MADM-11^TG/TG^* mice. To induce MADM clones, timed pregnant females were IP injected with TM (dissolved in corn oil; 1-2mg/mouse) at E12.5, E13.5, E14.5 and E15.5 respectively. At E19.5, litters were del ivered by caesarean section and raised with foster females. At P28, MADM-labeled animals were deeply anesthetized and fixed by transcardiac perfusion with 4% PFA, the brains were collected and postfixed overnight, cryoprotected in 30% succrose, embedded in Tissue-Tek O.C.T. (Sakura) stored at -20°C or -80°C until further use. Brains were cyosectioned at 50mm thickness, individual sections collected separately in PBS, and mounted onto superfrost glass slides (Thermo Fisher Scientific) keeping the same serial order and left-right orientation. Sections were counterstained with DAPI, air dried, coverslipped with Mowiol (475904-100GM, Merck) and analyzed. Serial sections were screened using an axioscope (Zeiss Axio Imager, Zeiss), and the presence of sparsely-labeled MADM cells was documented. Illustrative images were obtained with a Leica SPEII confocal microscope. MADM-labeled cells were considered to belong to a specific clone if they were within 300mm from their nearest neighbor in the tangential axis (x-z), typically found along consecutive sections. Clones were retained for analysis only if they were two-color clones - containing green and red neurons (clones with yellow cells were excluded), and if they were found within the neocortex. Clones with cells in cingulate, prefrontal, entorhinal or pyriform cortices were excluded. For each clone, the cortical area and the laminar location of each cell were recorded.

### Quantification and statistical analysis

#### Birthdating

Images of BrdU and DAPI labeling in the somatosensory cortex were obtained from each animal with an Olympus FV10 confocal microscope, and the average pixel intensity of BrdU signal along the cortical thickness was measured using Fiji. The cortical thickness was divided into 20 bins, and BrdU intensities from pixels within each bin were averaged. For each BrdU labeling stage, binned measures from independent animals were averaged. Plot curves were smoothed by neighbor bin averaging.

#### Mitotic cells

Cells were counted along the ventricular surface of the neocortex in 20mm-thick coronal sections. PH3+ cells located less than 3 cell diameters from the ventricular surface were considered apical, and those further away were considered basal mitoses; the number of subapical mitoses was residual. For each section the total cell count was normalized per length of VZ apical surface. To calculate the rate of neurogenesis from apical and basal mitoses, we considered the cell cycle exit rates in the developing mouse neocortex (Q) as defined in (Takahashi *et al*, 1996), and the relative abundance of apical or basal mitoses. If the proportion of basal mitoses was below Q (between E14.5 and E16.5), the neurogenic rate of basal mitoses was considered 100% (consensus for IPCs) and the remaining mitoses necessary to reach Q corresponded to apical neurogenesis. Given the definition of Q, Q=A*pNA+B*pNB, where A and B are the % of apical and basal mitoses, and pNA and pNB are the probability of apical and basal mitoses to make neurons, respectively, the rate of apical neurogenesis was calculated as pNA=(Q-B)/A, for pNB=1. Next, given pNA, pNB and the density of apical and basal mitoses at each age, we calculated the density of neuron-producing apical and basal mitoses at each age (source of neurons), and with this the frequency of apical and basal neurogenesis modes.

#### BrdU labeling

Images from a single confocal plane were obtained and analyzed using a Leica SPEII confocal microscope or Olympus FV10 confocal microscope. For cortical neuron birthdating analysis, GFP+ cells containing BrdU were quantified and classified according to their amount of BrdU label, as previously (Cárdenas *et al*, 2018): 100%, full nucleus; 50%, nucleus with clearly less BrdU label than 100% but more than 10 speckles; 25%, nucleus containing 10 speckles of BrdU or less (Extended Data Fig. 1c).

#### BrdU labeling correction

In BrdU + electroporation experiments, GFP+ cells with 100% BrdU signal were born by DN from Apical Progenitors (APs, or RGCs). Cells with 50% BrdU signal were born after an extra round of cell division, but this could be from IPCs (IDN) or from RGCs (DN). Hence, the abundance of cells with 50% BrdU or less is an overestimation of IDN. We assumed that IPs divide always once to produce two neurons, and that the measures from one embryonic day are valid to estimate the probability of DN of the daughter APs. Accordingly, we can correct the numbers of cells with 50% BrdU to account only for the indirectly-born neurons. Since every experiment labeled a different number of cells, we needed to assume that APs are dividing asymmetrically into one AP and another cell, which can be either IP or neuron (*number_APs_* = *number_directly-born neurons_* + *number_IPs_*). With this assumption we normalized the data from different ages and computed the correction. Given *d_t_* and *i_t_* the number of directly and indirectly generated neurons, respectively, at time *t* (in the case of indirectly, the IP is born at time *t*), we obtained

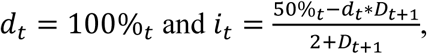

where *D*_t+1_ is the probability of an AP of producing a neuron instead of an IP at the time *t+1*. This correction requires an additional correction taking into account the next cell division, but at every step the magnitude of the correction decreases, so we considered the two extreme cases of *D*_t+2_ = 0 and *D*_t+2_ = 1 as the lower and upper bounds. We obtained two estimates of i, where *i_t_^min^* < *i_t_* < *i_t_^max^* and used them to calculate D:

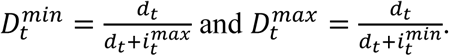

The final estimate of *D* is the average between the two extreme values, which are shown in the plot as the upper and lower bounds. The probability of IDN *I_t_* (production of an IP) is simply calculated as *I_t_* = 1 – *D_t_*.

#### MADM analysis of cortical clones

MADM-clonal analysis using MADM-11 transgenic lines has been previously validated to sparsely target single dividing progenitors giving rise to distinct clusters of cells each representing individual clonal units (Gao *et al*, 2014; Beattie *et al*, 2017). In particular, nearest-neighbor distance analysis was used to confirm clonality in sparse induction paradigm (Gao *et al*, 2014). Furthermore, high confidence of sparseness in labeling is attributed to the rare occurrence of MADM events in combination with a very low level of CreER activity (Beattie *et al*, 2020; Hippenmeyer *et al*, 2010; Zong *et al*, 2005). In the case when more than one cell cluster was found in the same brain, a 500mm minimum distance of separation was used as criteria to spatially distinguish separate clones. Data were statistically analyzed with Excel software using ANOVA, χ^2^-test, pair- wise *t*-test or independent samples *t*-test, where appropriate as indicated. Significance was set at *p* = 0.05. All values represent mean ± standard error of the mean (SEM).

## Notes

### Competing Interest Statement

The authors have declared no competing interest.

