## Supplemental Figures 1-4 for "Early Indirect Neurogenesis transitions to late Direct Neurogenesis in mouse cerebral cortex development"

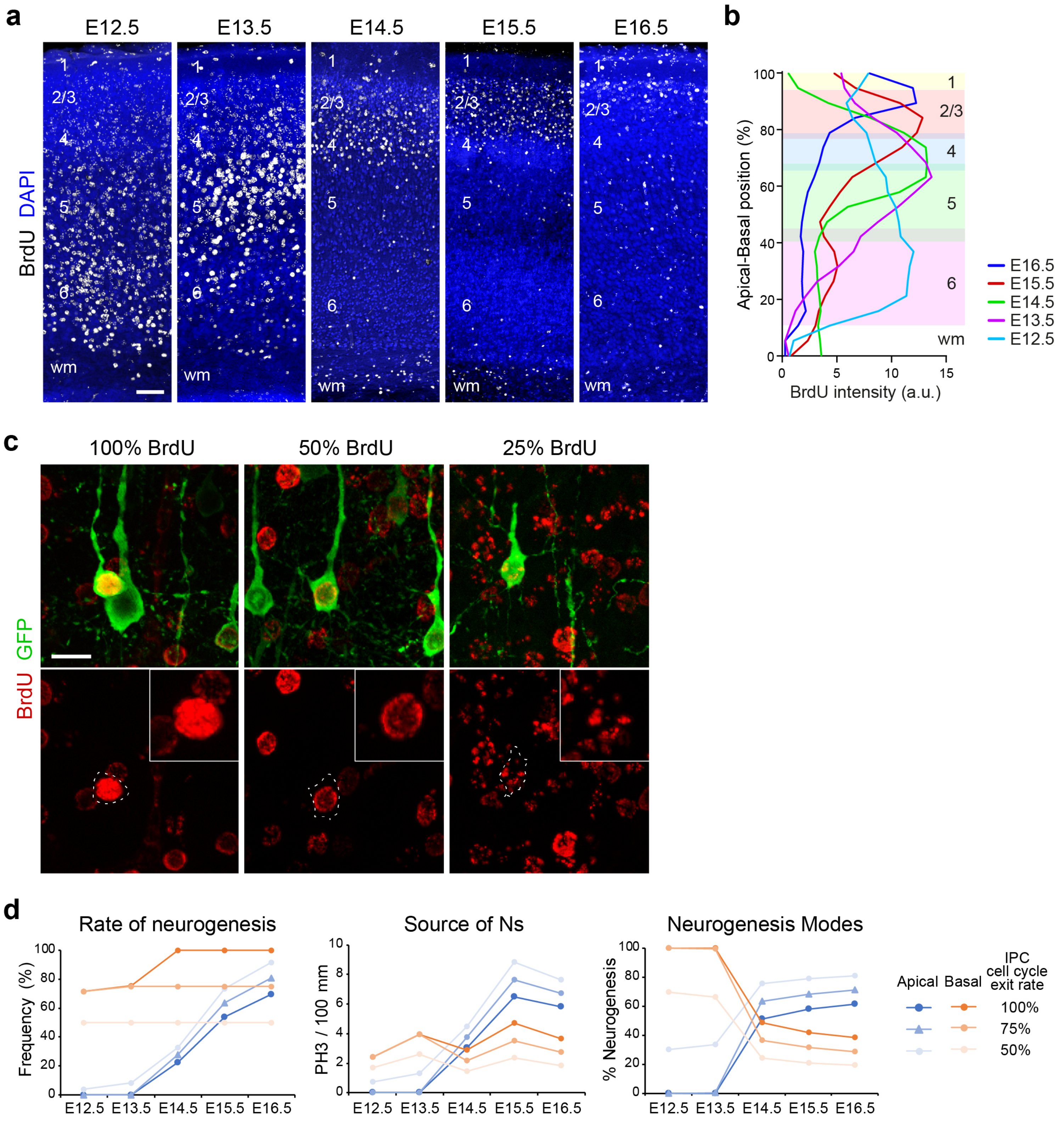


**Cárdenas, Extended Data Figure 1**

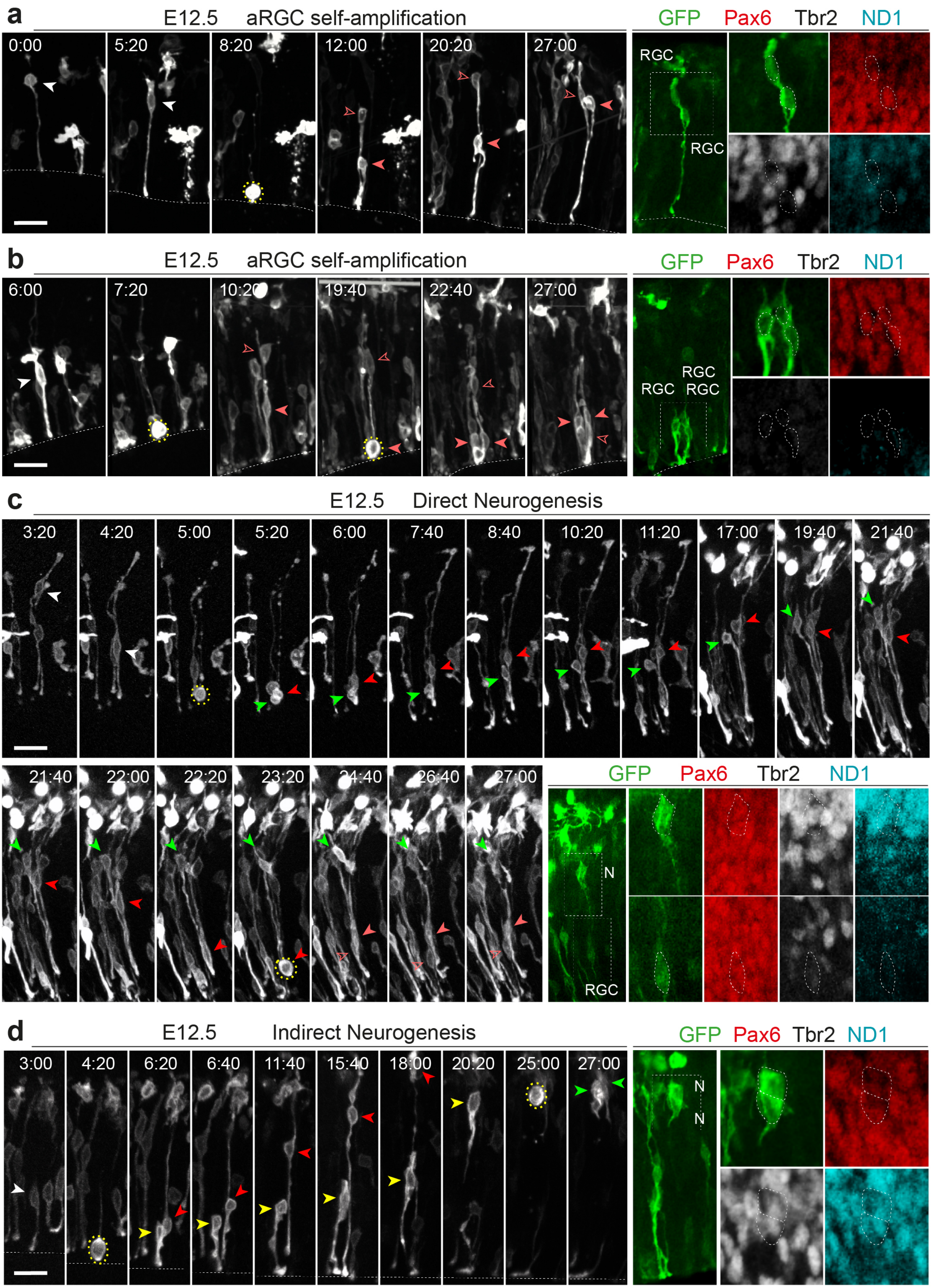


**Cárdenas, Extended Data Figure 2**

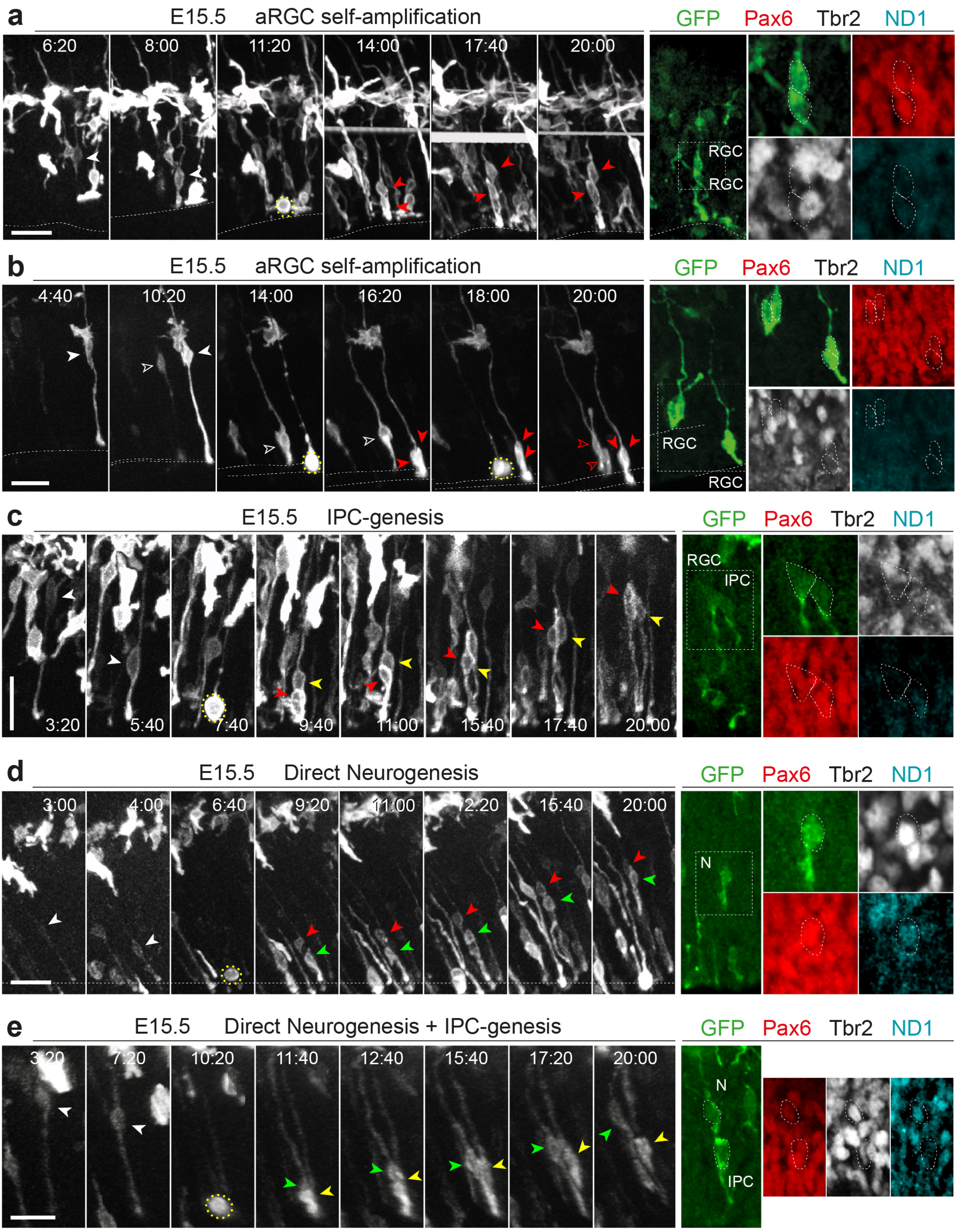


**Cárdenas, Extended Data Figure 3**

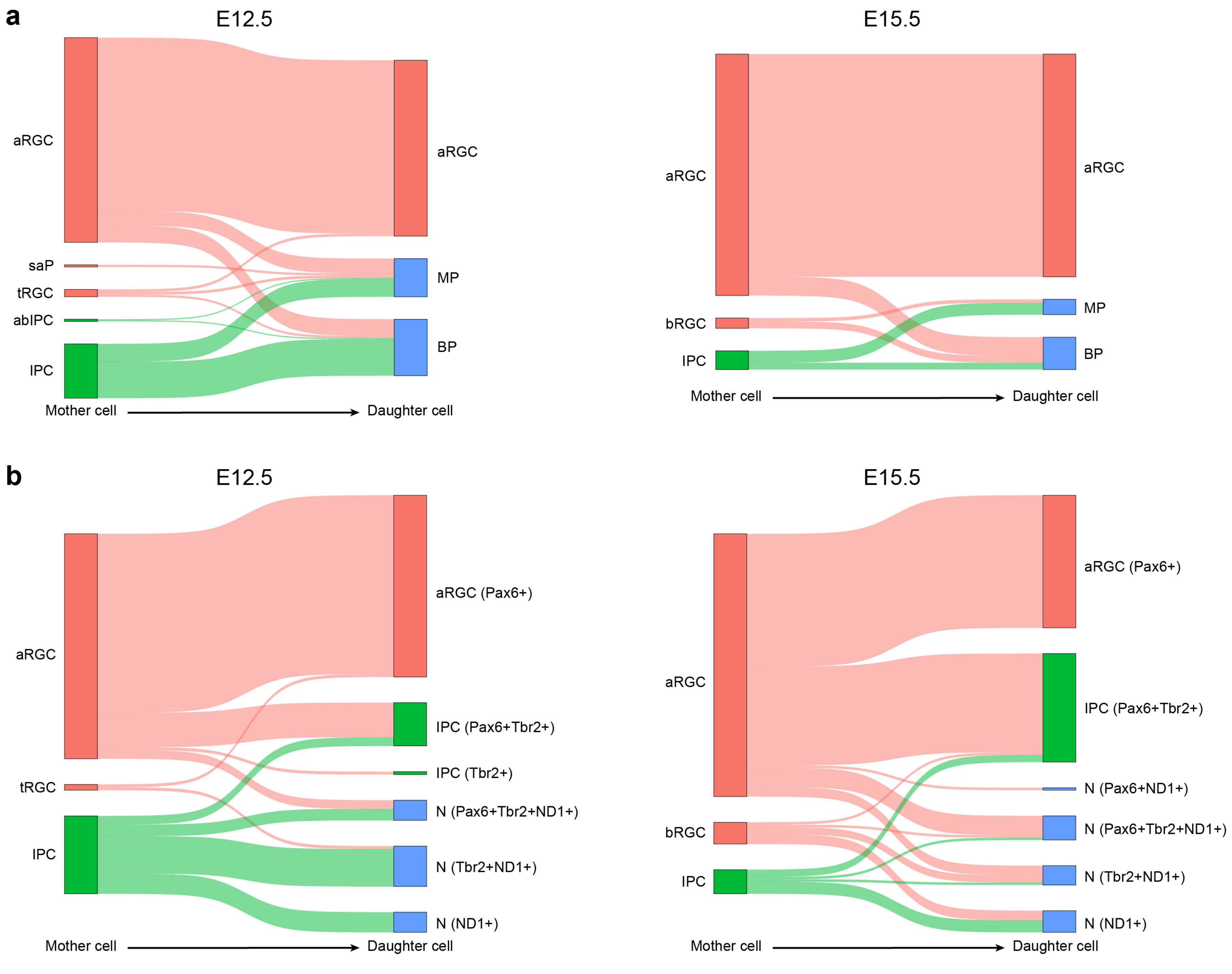


**Cárdenas, Extended Data Figure 4**
